# Shared and distinct responses of human and murine alveolar macrophages and monocyte-derived macrophages to *Mycobacterium tuberculosis*

**DOI:** 10.1101/2025.02.28.640814

**Authors:** Kimberly A Dill-McFarland, Glenna Peterson, Pamelia N. Lim, Shawn Skerrett, Thomas R Hawn, Alissa C. Rothchild, Monica Campo

## Abstract

Macrophages serve as important sites of bacterial replication and host immune response during *Mycobacterium tuberculosis* (Mtb) infection with distinct roles for alveolar macrophages (AMs) early in infection and monocyte-derived (MDMs) during later stages of disease. Here, we leverage data from human and mouse models to perform a cross-species analysis of macrophage responses to Mtb infection. Overall, we find that both subsets of human and murine macrophages mount a strong interferon response to Mtb infection. However, AM across both species do not generate as strong a pro-inflammatory response as human MDMs or murine bone marrow-derived macrophages (BMDMs), as characterized by TNFA signaling and inflammatory response pathways. Interestingly, AMs from mice that were previously vaccinated with BCG (scBCG) or from a model of contained TB (coMtb) had Mtb responses that were more similar to human AMs than control mice. We also identify species-specific pathways altered by infection differently in mouse and human macrophages, specifically in pathways related to cholesterol in AMs as well as MYC targets and Hedgehog signaling in MDMs/BMDMs. Lastly, to investigate downstream effects of the macrophage interferon responses, we examine macrophage expression of IL-10, an immunosuppressive cytokine induced by Type I Interferons, and c-Maf, a transcription factor required for IL-10 expression in myeloid cells. We find that c-Maf and IL-10 have significantly lower expression in AMs compared to MDMs in both humans and mice, suggesting one possible mechanism by which AMs mount a stronger interferon response following Mtb infection. Overall, these results highlight the dynamics of innate myeloid responses over the course of Mtb infection and the benefit of a combined analysis across species to reveal conserved and unique responses.

## INTRODUCTION

*Mycobacterium tuberculosis (*Mtb), the causative agent of tuberculosis (TB), led to 1.3 million deaths in 2022, making it one of the deadliest infectious agents worldwide (1). Macrophages are a critical replication niche for Mtb but also serve as an important site for the host innate immune response. Alveolar macrophages (AMs), the most abundant tissue-resident immune cell in a healthy lung, are the first cells productively infected with Mtb after aerosol transmission (2, 3). However, over the course of infection, there is an accumulation of recruited, rather than resident, myeloid cells that also serve as a replication niche for the bacteria (2–4). Interrogations of which myeloid populations serve as the major sites of bacterial replication identified that AMs provide an early permissive niche (5–7), while recruited monocyte-derived macrophages (MDMs) act as a site of bacterial replication later during infection (8, 9), especially after a CD4+ T cell response is mounted (10). Moreover, with advances in flow cytometry and single-cell sequencing, there is now greater appreciation for the phenotypic diversity of myeloid cells that can respond to lung infections (11). Our understanding of these different pulmonary myeloid populations is further complicated by fate-mapping and ontological studies that show changes in macrophage responses over time attributed to both changes in the dynamics of macrophage recruitment from the periphery as well as innate training of tissue-resident cells (12–15).

We recently found that human AMs from healthy volunteers mount a unique response to Mtb infection *in vitro* compared to paired human MDMs (16). Specifically, human AMs strongly up-regulated type I IFN genes, IFNA8 and IFNA1, while MDMs generated a robust response to Mtb through other pathways, including MYC and MTORC1. Similarly, in a murine study of *Mycobacterium* exposure, we found that murine AMs isolated from mice previously exposed to subcutaneous BCG vaccination (scBCG) or a model of contained TB infection (coMtb) mount a strong IFN-associated response to Mtb both *in vivo* and *in vitro* (*17*). This exposure-induced response is distinct from the response to Mtb generated by AM from naïve animals, which is characterized by up-regulation of antioxidant and anti-inflammatory genes and a dependence on the transcription factor NRF2 *in vivo*. Moreover, AM from naïve mice lack a robust pro-inflammatory response both *in vivo* and *in vitro* (*2*).

In addition to the studies described above, many others have documented transcriptional innate immune responses of murine or human AMs and MDMs (7, 18–20) as well as their metabolic changes in response to infection (5, 21, 22). However, very few studies have attempted to compare the early innate immune response to Mtb in tissue-resident AMs and recruited MDMs across human and murine samples. While animal models serve as a critical tool for studying host immune events during infection *in situ,* especially at early time points before most human infections are diagnosed, research animals cannot replicate every aspect of the human TB spectrum, especially the diversity of prior pulmonary exposures or respiratory diseases. Therefore, the opportunity to characterize macrophage responses across species is important for developing animal models that better replicate human disease, for assessing myeloid cells as novel therapeutic targets (23), and for understanding the range and specificity of macrophage immune responses to Mtb.

## RESULTS

### Comparing Macrophage Transcriptomes Across Human and Murine Species

We compared transcriptional responses to *Mycobacterium tuberculosis* (Mtb) infection across human and murine alveolar macrophages (AMs), human monocyte derived macrophages (MDMs), and murine bone marrow-derived macrophages (BMDMs) (**Figure 1A, B**). The human dataset consisted of 6 healthy donors’ paired AMs obtained by bronchoalveolar lavage (BAL) and 5-day differentiated MDMs isolated from peripheral blood mononuclear cells. Human macrophages were infected e*x vivo* for 6 hours with H37Rv (16). In human AM and MDM, there were 3921 and 8595 unique Mtb-induced differentially expressed genes (DEGs) at FDR < 0.1, respectively (**Table S1**). The murine dataset was composed of AMs from unvaccinated (control) C57BL/6J mice as well as mice vaccinated with subcutaneous BCG (scBCG) or exposed to a contained model of Mtb (coMtb). Murine AMs were obtained by BAL and then infected with H37Rv for 6 hours (17). Murine BMDMs were isolated from bone marrow of unvaccinated C57BL/6J mice and infected with H37Rv for 4 hours after 6 days of differentiation (22). Unique Mtb-induced murine macrophage DEGs included 1061 in control AMs, 2810 in scBCG vaccinated AMs, 4543 in coMtb AMs, and 6532 in BMDMs (FDR < 0.1, **Table S1**). When comparing Mtb-dependent DEGs between species (**Figure 1C**), we found 1664 orthologous DEGs by HGNC symbol significant in both human and at least one murine AM (15% of all AM DEGs). For MDMs, we found 3978 orthologous DEGs (35% of all MDM DEGs) were significantly impacted by Mtb infection in at least one treatment in both species.

**Figure 1.**
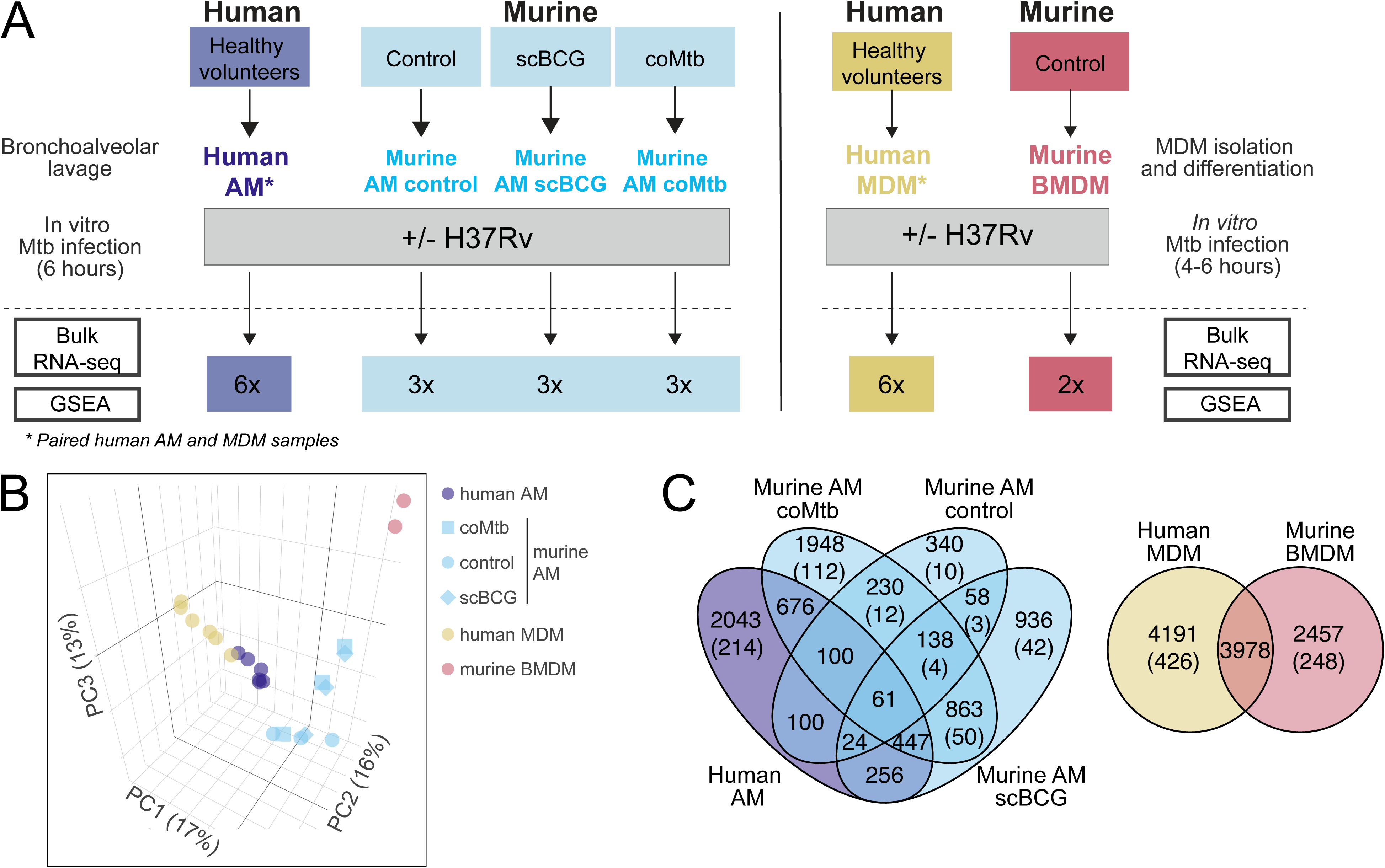
Study Design. (A) Diagram of human and mouse data study design and methods for data combination. (B) Principal component analysis (PCA) of human and murine macrophage gene expression in response to Mtb infection. Log_2_ fold change of +Mtb minus media in human and murine AMs and MDMs was calculated for genes expressed in both humans and mice. PCA was performed to visualize global gene expression with points colored by species and cell type. Shape indicates treatments within murine AM experiments. (C) Venn diagrams depicting Mtb-dependent differentially expressed genes (DEGs) in AMs and MDMs (FDR < 0.1). Values in parentheses indicate genes without an ortholog in the other species that are counted within the total. When multiple orthologs exist for murine MGI symbols, all equivalent human HGNC symbols are counted; thus, murine Venn totals may not represent unique genes within species and cell type.

### Shared Immune Pathways in Human and Murine Alveolar Macrophages Upon Mtb Infection

To examine species differences, we used Gene Set Enrichment Analysis (GSEA) using linear model estimates (Mtb vs media fold changes) against the MSigDB Hallmark database (**Figure 2A**, see Supplemental methods). GSEA was utilized to reduce potential batch effects found in DEG-level results through examination of pathways with orthologous genes and the use of average estimates without variance that may be attributed to experimental differences such as sample size and paired versus unpaired designs. GSEA revealed enrichment of 14 pathways in both human and murine AMs upon Mtb infection (FDR < 0.1, **Table S2**). The majority of these pathways are immune-related, such as TNFA signaling via NF-kB, which was consistently upregulated by Mtb infection in human and murine AMs regardless of exposure status (**Figure 2A**,). Similarly, other immune pathways including inflammatory, interferon gamma (IFNG), and interferon alpha (IFNA) responses were upregulated by Mtb infection in human and *Mycobacterium*-exposed murine AMs (scBCG, coMtb) (FDR < 0.1) but not control AMs (FDR > 0.1).

**Figure 2.**
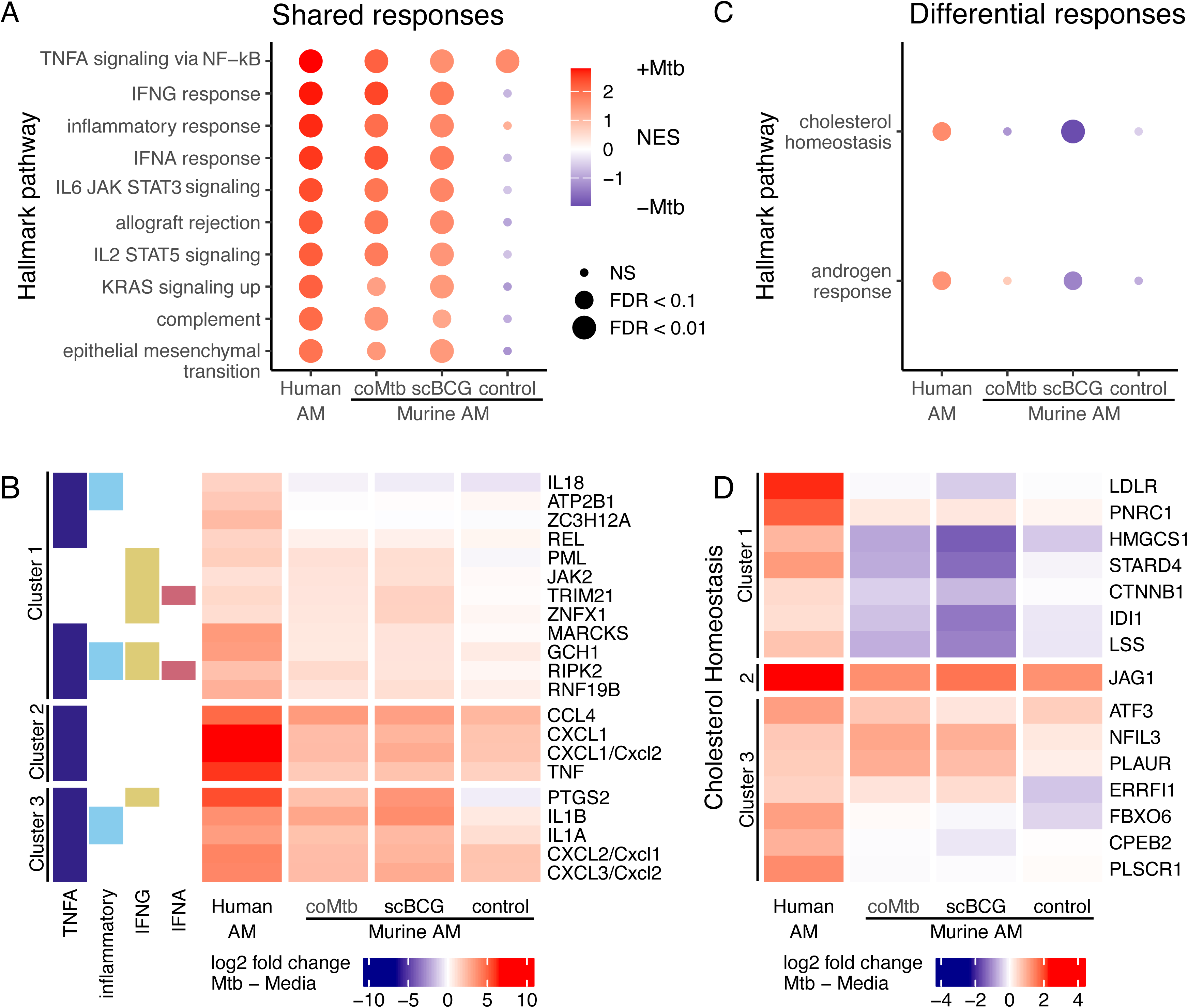
Shared and unique human and murine AM responses to Mtb infection. Gene set enrichment analysis (GSEA) of Mtb vs media gene expression in human and murine AMs. (A) Shared responses were defined as gene sets significantly enriched in the same direction in human and murine AMs (FDR < 0.1). The top gene sets are shown here (FDR < 0.01, N = 10 of 14) with full results in **Table S1**. (B) Genes were filtered to those expressed in both human and murine AMs, GSEA leading-edge in at least one selected pathway, and among the top 20 differentially expressed genes (DEGs) in any one AM (FDR < 0.01). Heatmap depicts log_2_ fold change of +Mtb minus media in human and murine AMs. Rows are clustered by Euclidean distance, and human HGNC symbols are used when human and murine gene names are identical. Left color annotation indicates the presence/absence of genes in each selected pathway including TNFA signaling via NK-kB (TNFA), inflammatory response (inflammatory), IFNG response (IFNG), and IFNA response (IFNA). (C) Unique responses were defined as gene sets significantly enriched in opposite directions in human and/or murine AM (FDR < 0.1). (D) Genes were filtered and heatmaps were clustered similar to (B) but for the cholesterol homeostasis pathway.

Gene-level analysis further supports these findings, including a heatmap of the leading-edge GSEA genes in selected pathways (TNFA, IFNG, inflammatory, IFNA in **Figure 2B**) that are expressed in both species. Consistent with GSEA directionality, many individual genes (Clusters 2 and 3) show elevated expression after Mtb infection in human AMs as well as murine coMtb and scBCG AMs. In particular, several chemokines (CCL4, CXCL1, CXCL2) are highly upregulated by Mtb infection with very low expression prior to infection (**Figure S1A**). The remaining top genes vary in baseline expression with the majority slightly more highly expressed in human compared to murine AMs. Overall, the data indicate robust activation of immune responses in both human and murine AMs following Mtb infection, with notable upregulation of IFNA and IFNG response pathways.

### Differential Activation of Immune Pathways in Mtb-infected Human and Murine Alveolar Macrophages

GSEA analysis reveals unique responses to Mtb infection between human and murine AMs (**Table S2, Figure 2C**). Most notably, the cholesterol homeostasis pathway is significantly upregulated in human AMs (normalized enrichment score [NES] = 1.6, FDR = 0.01), whereas murine scBCG AMs exhibit significant downregulation (NES = –2.0, FDR = 5E-5) of this pathway. Murine control and coMtb AMs show negligible changes (FDR = 0.5, 1). Cluster 1 highlights cholesterol genes such as HMGCS1, IDI1, and LSS that were upregulated in human AMs and generally downregulated in murine AMs regardless of vaccination status (**Figure 2D**, **Figure S1B**). This may indicate a decrease in cholesterol biosynthesis after Mtb infection in murine AMs that is not observed in human AMs. In contrast, PLSCR1 (Cluster 3) and LDLR (Cluster 1) display patterns of upregulation in human AMs but minimal changes in murine AMs. This may indicate differences in lipid transport in the two species after Mtb infection.

While IFNG and IFNA response pathways were consistently upregulated in AMs in response to Mtb (**Figure 2A**), a targeted analysis of IFN genes and receptors revealed differences between the species (**Figure S2, Table S1**). Specifically, human AMs showed robust upregulation of several Type I, II, and III IFNs (IFNA1, IFNA8, IFNA13, IFNB1, IFNG, IFNL1) after Mtb infection while murine AMs showed minimal changes and often no detectable expression of IFNs. Additionally, androgen response was downregulated in murine AMs in response to Mtb infection compared to human AMs (**Figure 2C**). The androgen response pathway may be implicated here due to bias in the data sets with inclusion of both male and female human donors but only female mice. However, further research is needed to investigate these possible divergent hormone responses in human AMs compared to murine AMs in response to early Mtb infection.

### Species-Specific Transcriptional Responses to Mtb Infection in Human and Murine MDMs

Overall, both human MDM and murine BMDM mount strong innate responses to Mtb infection, with significant enrichment for pro-inflammatory pathways such as TNFA signaling via NF-kB, inflammatory response, and IL6 JAK STAT3 signaling as well as IFNG response (**Figure 3A**, **Table S2**). At the individual gene level, many genes were equivalently expressed by both human MDM and murine BMDM (Clusters 1 and 4), while others were more strongly expressed by murine BMDM (Clusters 2 and 3), most notably CXCL11 (**Figure 3B, Figure S3A**). Similar to AMs, IFN production itself varied between the species (**Figure S2**, **Table S1**). Human MDMs appear to preferentially upregulate IFNB1 and IFNG with no detection of IFNAs, while murine BMDMs upregulate several IFNAs as well as IFNB1 in response to Mtb.

**Figure 3.**
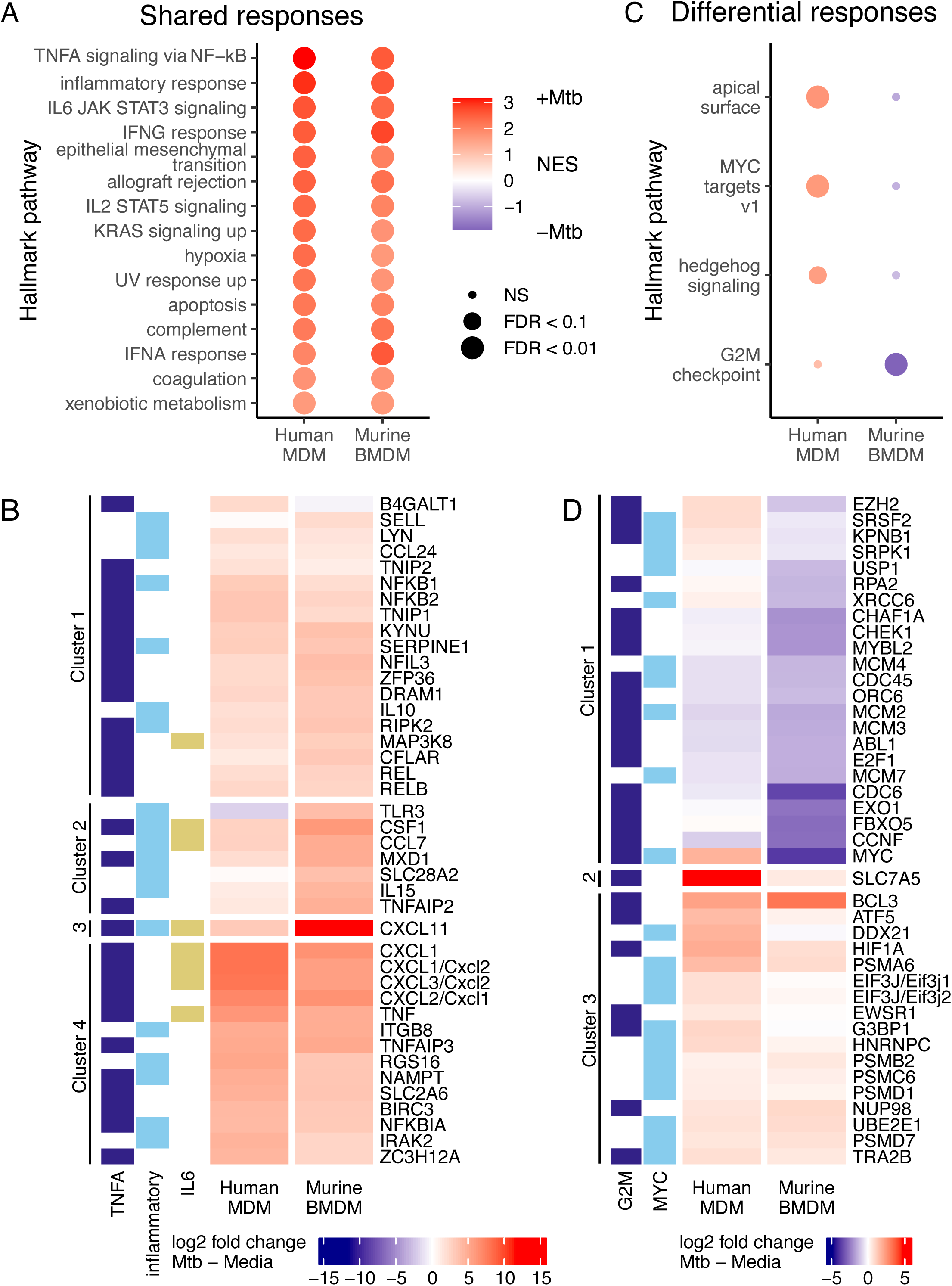
Shared and unique human and murine MDM responses to Mtb infection. Gene set enrichment analysis (GSEA) of Mtb vs media gene expression in human and murine MDMs. (A) Shared responses were defined as gene sets significantly enriched in the same direction in human and murine MDM (FDR < 0.1). The top gene sets are shown here (FDR < 0.01, N = 14 of 15) with full results in **Table S1**. (B) Genes were filtered to those expressed in both human and murine MDM, GSEA leading-edge in at least one selected pathway, and among the top 20 differentially expressed genes (DEGs) in either MDM (FDR < 0.01). Heatmap depicts log_2_ fold change of +Mtb minus media in human and murine MDMs. Rows are clustered by Euclidean distance, and human HGNC symbols are used when human and murine gene names are identical. Left color annotation indicates the presence/absence of genes in each selected pathway including TNFA signaling via NK-kB (TNFA), inflammatory response (inflammatory), and IL6 JAK STAT3 signaling (IL6). (C) Unique responses were defined as gene sets significantly enriched in opposite directions in human and/or murine MDM (FDR < 0.1). (D) Genes were filtered and heatmaps were clustered similar to (B) but for G2M checkpoint (G2M) and MYC targets v1 (MYC) pathways.

Gene sets were filtered to those significant in either human MDM or murine BMDM (FDR < 0.1) as well as enriched in different directions in human and murine MDMs. This analysis revealed no gene sets significantly upregulated in opposite directions in human vs murine MDMs (**Figure 3C, Table S2**), but 4 pathways had opposing trends. Specifically, the apical surface, MYC targets v1, and Hedgehog signaling pathways were significantly upregulated in human MDMs upon Mtb infection (FDR < 0.1) and non-significantly downregulated in murine MDMs (FDR > 0.1). In addition, the G2M checkpoint pathway showed the opposite trends. Leading-edge DEGs confirmed GSEA directionality with Cluster 1 (notably MYC) showing genes more strongly downregulated in murine BMDM than human MDM and Clusters 2 and 3 showing stronger upregulation in human MDM. (**Figure 3D**). Overall, these pathway genes have moderate expression in media and moderately higher expression after Mtb infection (**Figure S3B**). These differential gene expression profiles underscore species-specific responses to Mtb infection by MDMs emphasizing the importance of context-specific analysis in understanding host-pathogen interactions.

### Macrophage Subset-Specific Responses to Mtb Shared Across Species

To define gene expression pathways that were specific to each macrophage subset, but shared across species, gene sets were filtered for those significant in the same direction in both species in either uninfected or Mtb-infected samples (FDR < 0.1, **Figure 4**). Directly comparing gene expression of AM to MDM within species, we found that uninfected human and murine MDMs are both enriched for E2F targets and G2M checkpoint pathways compared to AMs, while AMs are enriched for TNFA signaling via NF-kB and myogenesis pathways compared to MDMs (**Figure 4**, **Table S2**). For Mtb infected samples, three pathways are significantly enriched in MDMs compared to AMs, which were not enriched at baseline: TNFA signaling via NF-kB, inflammatory response, and allograft rejection. These findings highlight functional differences between MDMs and AMs that shift with infection, with AMs expressing a more inflammatory profile than MDMs at baseline, but MDMs mounting a stronger pro-inflammatory response following Mtb infection compared to AMs (**Figure 4**). These data reveal that some major differences between responses of tissue-resident macrophages and recruited macrophages are shared across species.

**Figure 4.**
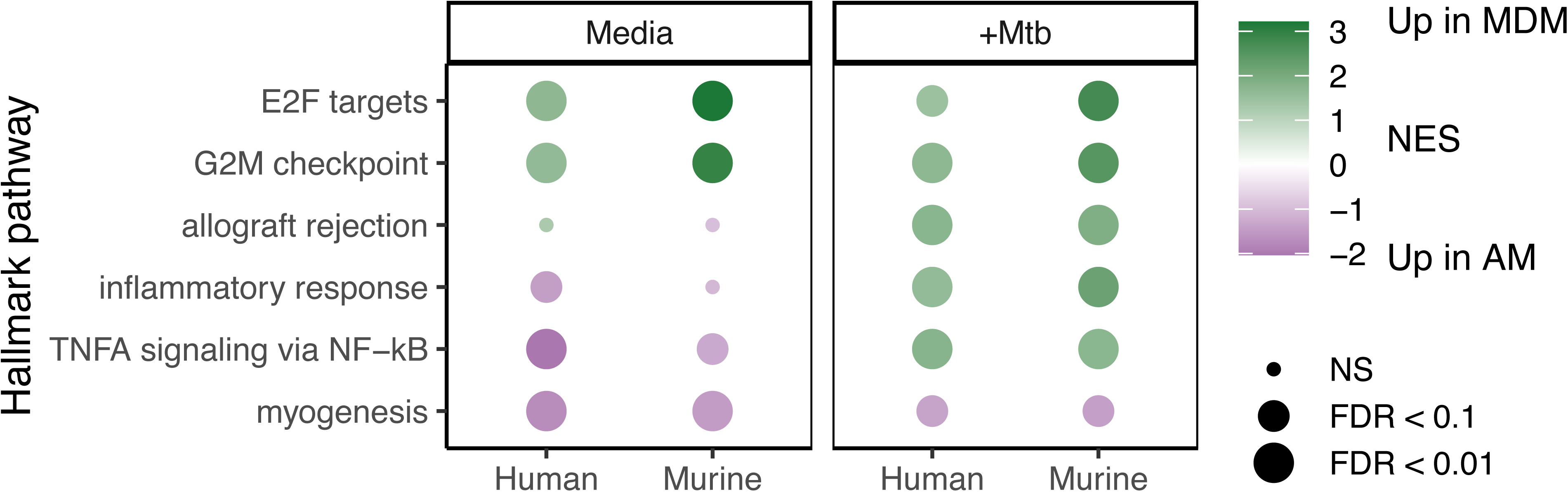
Differential enrichment of hallmark pathways in MDMs vs AMs in response to Mtb infection. Gene set enrichment analysis (GSEA) of AMs vs MDMs in media (left) and Mtb-infected (right) conditions. For mice, control AMs were used in comparisons. Gene sets were filtered to those significant in the same direction in both species in at least 1 infection condition (FDR < 0.1). Normalized enrichment scores (NES) with green indicating upregulation in MDMs and purple indicating upregulation in AMs.

### Candidate Gene Analysis Finds Differential Expression of MAF and IL10 in Alveolar and Monocyte-Derived Macrophages Upon Mtb Infection

Following our pathway analysis approach, we took a candidate gene approach to examine expression of specific genes and pathways that have known roles during Mtb infection (24–26) (**Figure S4**). We examined genes associated with macrophage antimicrobial responses (CAMP, IL1B, NOS1, NOS2, TNF, LYZ/Lyz2), genes with known Mendelian Susceptibility to Mycobacterial infection (GATA2, IRF8, ISG15, STAT1) as well as genes involved in Reactive Oxygen Species or other innate pathways (CYBB, CD14, HIF1A, NLRP3). As previously reported, we found major differences in the expression of NOS2 across species, with the gene undetectable in all human samples (27, 28).

Prior studies have found that expression of c-MAF and MAFB distinguish tissue-resident AM from recruited macrophage subsets in the lung and that low expression of c-MAF and MAFB are critical for the ability of tissue-resident macrophages to undergo self-renewal (29, 30). Additionally, our previous study found that murine AMs lack of IL-10 production is due to low expression of the transcription factor c-MAF, while murine BMDMs express high levels of c-MAF and are able to produce IL-10 (31). To determine whether MAF, the gene that encodes for the protein c-MAF, and IL10 gene expression differences between AM and MDM were shared across human and murine macrophages following Mtb infection, we examined the expression profiles of MAF (**Figure 5A**) and IL10 (**Figure 5B**) in human AMs and MDMs under media and Mtb-infected conditions (**Table S1**). In uninfected cells, MAF expression was significantly higher in MDMs than AMs for both human (FDR = 6E-21) and murine cells regardless of exposure status (FDR < 3E-10, **Figure 5A**). Similarly, IL10 expression was also significantly higher in uninfected human (FDR = 7E-6) and murine MDMs (FDR < 1E-6) compared to AMs (**Figure 5B**).

**Figure 5.**
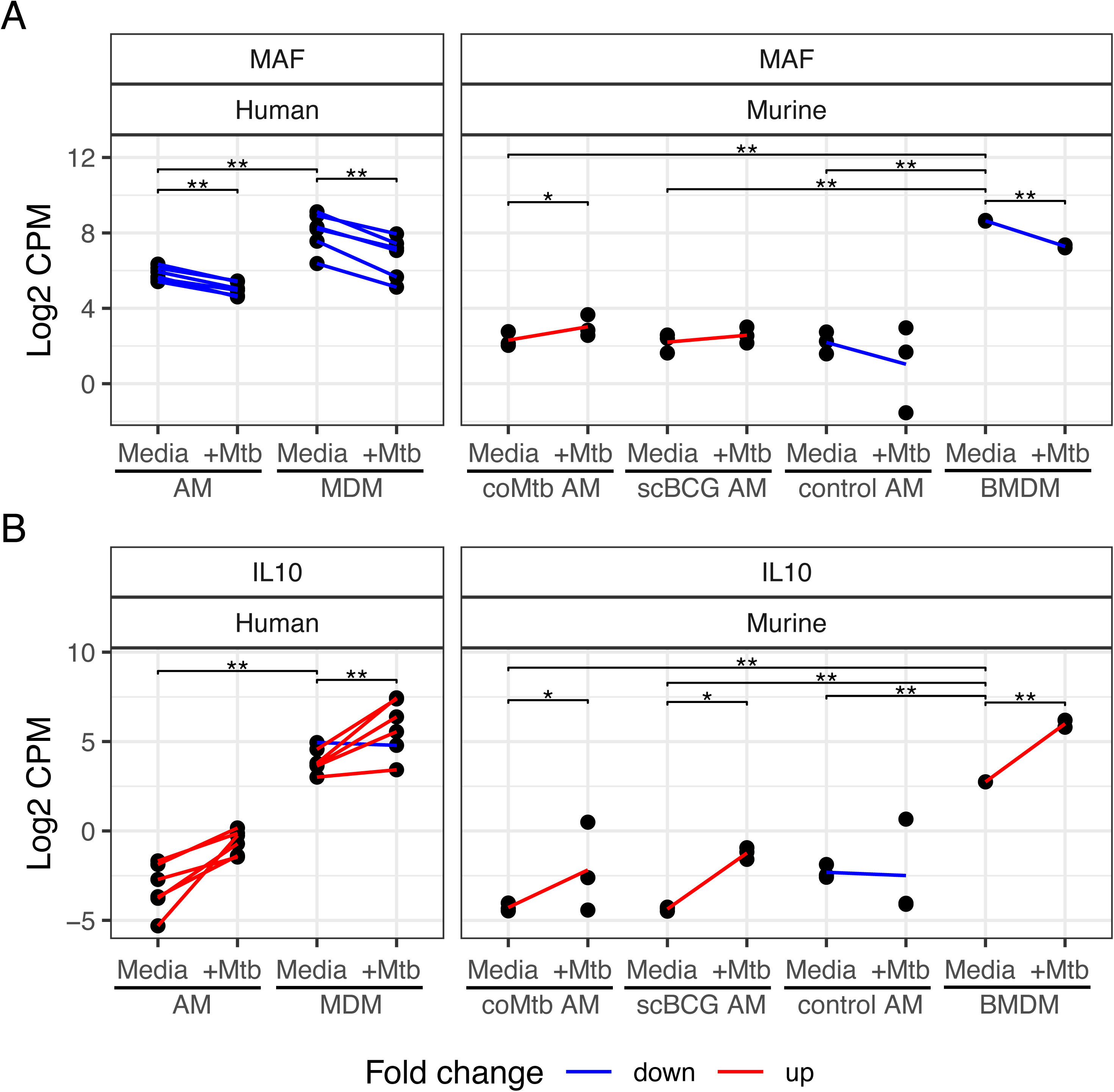
MAF and MAF-regulated IL10 expression as markers of AM. Log_2_ normalized counts per million (CPM) expression of (A) MAF and (B) IL10 gene expression in human and murine AMs and B/MDMs. Paired media and +Mtb samples are connected by a line for human samples while a single line indicates the difference in mean media and +Mtb expression for unpaired murine samples. These lines are colored by positive (red) vs negative (blue) fold change. Significance is indicated for Mtb infection within each cell type and between cell types in the media condition. ** FDR < 0.01; * FDR < 0.1

We also observed effects of Mtb infection on MAF and IL10 gene expression. MAF was significantly downregulated in human AMs and MDMs as well as murine BMDMs (FDR < 0.01) following Mtb infection. In contrast, IL10 was significantly upregulated in response to Mtb in all B/MDMs and AMs except murine control AMs. These findings underscore the distinct regulatory mechanisms and immune responses of AM and MDM to Mtb infection in humans and mice. The changes in gene expression following Mtb infection also suggest that while MAF appears to be required for IL10 expression, there are likely additional modulatory factors.

## DISCUSSION

Although the fundamental pathways governing early inflammation and immune defense are well conserved across species, significant differences exist in how human and murine macrophages respond to Mtb at the transcriptional level. Previous studies have established that TNFA, IFNG, and other pro-inflammatory pathways are critical in controlling Mtb infection (24, 32–35), yet the specific dynamics within AMs and MDMs across species have remained less explored. In this study, we compared how human and murine AMs and MDMs respond to early Mtb infection at the transcriptional level, uncovering both shared and distinct immune responses. There was a high degree of conservation in immune responses between human and murine AMs following Mtb infection. Both Mtb-infected human AMs and murine AMs from scBCG vaccinated mice and coMtb-exposed mice upregulated genes from key inflammatory pathways such as TNFA signaling via NF-κB, IFNG response, inflammatory response, and IFNA response. These shared pathways underscore the fundamental role of AMs as early responders to Mtb, where they must initiate crucial responses that can shape the trajectory of infection in both species. In particular, the upregulation of IFNA responses is consistent with previous studies identifying Type I IFN cytokines as critical mediators of macrophage activation and bacterial susceptibility during Mtb infection in both humans and mice (36–39). Interestingly, murine AMs from control mice showed significantly lower enrichment for many of these inflammatory pathways compared to all of the other groups. This may indicate that prior infection and trained immunity play a significant role in early Mtb immune responses (40).

In contrast, our data revealed that human and murine AMs manage cholesterol metabolism differently during Mtb infection *in vitro*. Human AMs showed a notable enrichment in the cholesterol homeostasis pathway after infection, whereas murine AMs had decreased enrichment in the pathway. Cholesterol is a preferred carbon source for Mtb (41, 42) and cholesterol metabolism contributes to macrophage responses to infection, such as the development of foamy macrophages in response to Mtb (41, 43–45). Transcriptional differences in cholesterol and lipid metabolism have also been implicated in human infection phenotypes (46). Thus, differences in human and murine cholesterol pathways in response to Mtb infection may influence disease progression *in vivo*, making it an important area of further research.

One of the key differences between tissue-resident AMs and recruited macrophages lies in how they regulate inflammation during Mtb infection. Our analysis found that while both AMs and MDMs upregulated genes from interferon and inflammatory pathways in response to Mtb in both humans and mice, MDMs mounted a more robust response in many genes from pro-inflammatory pathways. This was further highlighted by differential enrichment of hallmark pathways in MDMs and AMs, which identified TNFA signaling via NF-kB, inflammatory response, and allograft rejection as the pathways most significantly enriched in Mtb-infected MDMs compared to AMs. This gene expression analysis supports functional data that MDMs are initially more capable of controlling intracellular bacterial replication compared to AMs *in vivo* (5, 7, 10).

Through candidate analysis, we observed that AMs and MDMs differed in their expression of MAF and IL10 with human and murine tissue-resident AMs exhibiting lower levels of MAF and IL10 compared to MDMs, which is consistent with studies showing c-MAF is a transcription factor that is associated with differentiation of monocytes to macrophages (47). Since c-MAF binds directly to the IL-10 gene promoter and is essential for driving IL-10 production (48), a powerful anti-inflammatory cytokine, the reduced levels of c-MAF and IL-10 in resident AMs suggest that these cells lack a critical autocrine brake following external activating signals and may respond more strongly. On the other hand, during later stages of Mtb infection, MDMs are recruited to the lung (5) and display high levels of both c-MAF and IL-10, which can benefit the host by preventing tissue damage but can also create a more permissive environment for Mtb. These data support growing evidence that, despite robust Mtb control by MDMs during early infection, Mtb-infected MDMs are less responsive than AMs to T cell-derived signals during chronic stages of infection (9, 10). In fact, a direct link between c-MAF, IL-10, and Mtb replication was previously identified in MDM cells in Dhiman et al (49). The authors showed that Mtb grows faster in CD14(hi) than in CD14(lo)CD16(+) cells and attributed it to cells having increased expression of c-MAF, which induced more IL-10 and hyaluronan. However, the overall decrease in MAF expression yet increase in IL10 expression after Mtb infection found here highlights that there are additional factors, separate from MAF, that also regulate IL10 expression during infection.

There are several limitations to our study. All of the gene expression analyses were performed on bulk macrophage samples, which ignores heterogeneity within each population. Both our group and others have found unique sub-populations within the tissue-resident AM population in both murine and human lungs (7, 17, 50). Similar findings of heterogeneity have been identified for human bone marrow and murine BMDM (9, 51). This heterogeneity could not be assessed in our bulk experiments. Additionally, our human samples and animal models do not fully reflect the known spectrum of TB disease and exposure. The human macrophage samples were all collected from healthy adults with minimal TB exposure. The murine studies were all performed in C57BL/6J mice and all studies used the H37Rv strain of Mtb. Additional studies across a diversity of mouse and Mtb strains will be needed to investigate how much host and pathogen diversity impacts macrophage responses. The scBCG and coMtb murine models provide some insight into the plasticity of the AM population following prior *Mycobacterium* exposure. However, the durability of the remodeling and the specific cellular signals that initially drive the changes in AM responses has not yet been addressed. Lastly, macrophage responses are significantly different *in vivo* compared to *in vitro*, where they receive many different signals from the lung environment (14, 52). We have previously found that Mtb-infected AMs generate a cell-protective, NRF2-dependent response, which is not observed *in vitro* (2). These studies were performed exclusively *in vitro* and therefore, do not incorporate lung-specific effects.

Overall, our results reveal both the conserved and unique Mtb responses of different macrophage types across species. Our study also highlights the need for more human studies across a spectrum of TB exposure and disease as well as new animal models to study the most critical pathways that shape macrophage immune responses to Mtb. These findings demonstrate the complex, yet mostly conserved, nature of immune defenses across species, while also pointing to important differences that influence how each species responds to Mtb.

## METHODS

### Data cleaning

We analyzed transcriptional responses to *Mycobacterium tuberculosis* (Mtb) infection across human and murine alveolar macrophages (AMs), as well as monocyte-derived macrophages (MDMs) and bone marrow-derived macrophages (BMDMs) from previously aligned and quantified gene expression tables (16) (2) (17). The human dataset included paired AM and MDM with and without *in vitro* Mtb infection from 6 healthy donors (N = 24 samples). The murine dataset included AMs and BMDMs from naïve (N = 3 AM, 2 BMDM), scBCG (N = 3 AM), and coMtb (N = 3 AM) mice with and without *in vitro* Mtb infection (N = 22 samples) (**Figure 1A**). In order to harmonize methods, all datasets were processed from raw counts using the same pipeline in R v4.3.1 (53). Human and murine datasets were cleaned and modeled independently. Raw counts were filtered to protein-coding genes with at least 0.1 counts per million (CPM) in at least 2 samples per dataset. Counts were trimmed mean of means (TMM) normalized and transformed to log2 CPM with gene-level quality weights with limma (54, 55). This resulted in 16,201 genes in the human dataset and 12,874 genes in the murine dataset. Principal component analysis (PCA) was performed using the 12,036 genes expressed in both AMs and MDMs with corresponding human and murine gene symbols. Log2 fold change was calculated for Mtb minus media, and PCA was performed on fold changes with scaling and centering.

### Gene-level linear models

Gene expression was modeled for Mtb infection and macrophage type with an interaction model in kimma (56). For human samples, cells included MDMs and AMs and a random effect was included to account for paired donors, ∼cell+Mtb+cell:Mtb + 1|donor. For murine samples, cells included BMDM, control AM, coMtb AM, and scBCG AM and a simple linear model was used, ∼cell+Mtb+cell:Mtb. All models also utilized gene-level weights. Differentially expressed genes (DEGs) were defined at FDR < 0.1.

### Gene set enrichment analysis (GSEA)

Mtb-infected vs media gene expression log2 fold changes were extracted from kimma contrast model estimates. Fast GSEA (57) was performed using Broad MSigDB Hallmark gene sets (58, 59). Significant pathways were defined at FDR < 0.1. In AMs, unique human or murine pathways were defined as those significant in human and at least one murine cell type (naïve, coMtb, scBCG) but enriched in opposite directions between species. In MDMs, unique pathways were defined similarly except significant was only required in one human or murine cell type due to lack of overlap. Shared human and murine pathways were defined as those that were significant and enriched in the same direction in human and at least one murine cell type (coMtb, scBCG). This stricter definition was applied to shared responses due to the high overlap of signal. In heatmaps, shared pathways were further restricted to FDR < 0.01, and genes were filtered to those expressed in both human and murine cells, GSEA leading-edge in at least one selected pathway, and among the top 20 differentially expressed genes (DEGs) across all pathways and cell populations (FDR < 0.01).

### Methods for prior murine studies

#### Mice

All mouse studies were previously described (17, 22). No new animal studies were performed for this manuscript. All previous animal studies were performed in compliance with and approval by the Seattle Children’s Research Institute’s Institutional Animal Care and Use Committee. All mice were housed and maintained in specific pathogen-free conditions. C57BL/6 mice were purchased from Jackson Laboratories (Bar Harbor, ME). 6-12 week old female mice were used for RNA-sequencing. Mice infected with Mtb were housed in Biosafety Level 3 facilities in Animal Biohazard Containment Suites.

### Murine *Mycobacterium* exposure models: BCG subcutaneous immunization (scBCG) and establishment of contained TB infection (coMtb)

Mice with scBCG and coMtb exposures were prepared as previously described (17). Briefly, BCG-Pasteur was cultured in Middlebrook 7H9 broth at 37°C to an OD of 0.1–0.3. Bacteria was diluted in PBS and 1 x 10^6^ CFU in 200 ml was injected via subcutaneous route (scBCG). Intradermal infections to establish coMtb were performed by injecting 10,000 CFU of Mtb (H37Rv) in logarithmic phase growth intradermally into the ear in 10 μL PBS using a 10 μL Hamilton Syringe (coMtb), following anesthesia with ketamine/xylazine.

### Murine Alveolar Macrophage Isolation

AMs were isolated by BAL and pooled from 5 mice per group as previously described (2, 60). Briefly, the trachea of euthanized mice was exposed and then punctured with Vannas Micro Scissors (VWR). 1 mL PBS was injected using a 20G-1” IV catheter (McKesson) connected to a 1 mL syringe. The PBS was flushed into the lung and then aspirated three times and the recovered fluid was placed in a 15mL tube on ice. The wash was repeated 3 additional times. Cells were filtered and spun down. Cells were plated at a density of 5 x 10^4^ cells/well (96-well plate) in complete RPMI (RPMI plus FBS (10%, VWR), L-glutamine (2mM, Invitrogen), and Penicillin-Streptomycin (100 U/ml; Invitrogen) and allowed to adhere overnight in a 37°C humidified incubator (5% CO_2_). Media with antibiotics and non-adherent cells were washed out prior to stimulation.

### Murine Bone Marrow Derived Macrophages Isolation

As previously described (22), bone marrow was harvested from the femur and tibia by and homogenized by filtering through a 70 μm filter before being placed at 37 C, 5% CO2 in four 15 cm plates and cultured in BMDM media (RPMI 1640 supplemented with recombinant M-CSF ( 50 ng/ml), L-glutamine (4mM) and 10% fetal bovine serum, penicillin and streptomycin. Fresh BMDM media was added on day 3 or 4 and then on day 6, the adherent cells were washed twice and lifted with PBS + 2mM EDTA. Cells were replated in BMDM media without antibiotics at 400,000 cells/well in a 24-well plate and rested overnight before infection.

### Murine AM and BMDM Mtb Infection

H37Rv was prepared by culturing from frozen stock in 7H9 media at 37°C for 48 hours to O.D. of 0.1-0.3. The final concentration was calculated based on strain titer and bacteria was added to macrophages for two hours. Cultures were then washed three times to remove extracellular bacteria. Cell cultures were washed once in PBS after 6 hours (AMs) or 4 hours (BMDMs) to remove dead cells and collected in TRIzol for RNA isolation.

### Murine RNA Sequencing and Data Processing

For AMs, RNA isolation was performed using TRIzol (Invitrogen), two sequential chloroform extractions, Glycoblue carrier (Thermo Fisher), isopropanol precipitation, and washes with 75% ethanol. RNA was quantified with the Bioanalyzer RNA 6000 Pico Kit (Agilent). cDNA libraries were constructed using the SMARTer Stranded Total RNA-Seq Kit (v2) – Pico Input Mammalian (Clontech) following the manufacturer’s instructions. Libraries were amplified and then sequenced on an Illumina NextSeq. For BMDMs, total RNA was extracted using Direct-zol-96 extraction kits, including an on-column DNase treatment. RNA integrity was confirmed by Nanodrop One Microvolume UV-Vis Spectrophotometer. Illumina TruSeq stranded kits were used for library prep and libraries were sequenced on an Illumina NovaSeq6000 S4, producing 100 bp paired end reads. Read pairs were aligned to the mouse genome (mm10) + Mtb H37Rv genome using the gsnap aligner (v. 2018-07-04) allowing for novel splicing. Genes with low expression were filtered using the “filterByExpr” function in the edgeR package (61).

### Methods for prior human studies

#### Human Subjects

All human studies were previously described (16). Six healthy volunteers underwent phlebotomy to collect PBMCs and serum, as well as fiberoptic bronchoscopy to obtain alveolar macrophages via bronchoalveolar lavage. The study protocols were approved by the University of Washington’s Human Subjects Review Board, and all participants provided written informed consent.

### Human Alveolar Macrophage Isolation

AMs were collected from the same healthy, non-smoking volunteers (aged 18-50) without lung disease. AMs were obtained through fiberoptic bronchoscopy, with bronchoalveolar lavage fluid filtered, centrifuged at 300g for 10 minutes at 4°C, and washed in HBSS (without calcium or magnesium). Cells were resuspended in RPMI with 10% heat-inactivated FBS, 1% HEPES, 2mM L-glutamine, 100 U/mL penicillin, and 100 µg/mL streptomycin. Cell viability was confirmed via trypan blue exclusion, and macrophage purity was verified with Giemsa staining. AMs were then plated at 1E6 cells per well, washed, and cultured overnight in RPMI with 20% autologous serum, with infections occurring 24 hours post-harvest.

### Human Monocyte-Derived Macrophages Isolation

Peripheral blood mononuclear cells (PBMCs) were collected from the same healthy, non-smoking volunteers as for AMs. Blood samples were processed using Vacutainer CPT Cell Preparation Tubes (BD cat no. 362753), while PBMCs for experiments involving monocyte-derived macrophages (MDMs) only were obtained from leukoreduction chambers (Bloodworks Northwest, Seattle, WA). PBMCs were isolated via Ficoll gradient separation, cryopreserved, and stored in liquid nitrogen until use. For experiments, PBMCs (40E6) were thawed in batches and differentiated over five days in RPMI/10% FBS with 50 ng/mL macrophage-colony stimulating factor (M-CSF) to generate MDMs, which were then purified through magnetic bead separation (CD14 negative isolation protocol, Monocyte Isolation Kit II, Miltenyi Biotec). CD14+ MDMs were plated at 2E6 cells per well in RPMI with 20% autologous serum and maintained overnight at 37°C and 5% CO₂ before infection or stimulation.

### Human AM and MDM Mtb Infection

Thawed H37Rv (ATCCR 25618TM) stock cultures were grown in 7H9 medium supplemented with ADC enrichment, 0.05% Tween 80, and 0.2% glycerol in a shaking incubator at 37°C and 150 RPM. Cultures were washed, resuspended in Sauton’s medium, and stored as stocks. For infection, macrophages were exposed to these thawed Mtb stocks at a multiplicity of infection (MOI) of 1:1. Following a 6-hour incubation at 37°C with 5% CO₂, cells were lysed in TRIzol and preserved at –80°C.

### Human RNA Sequencing and Data Processing

RNA extraction included thawing TRIzol samples, chloroform phase separation, and ethanol precipitation, followed by processing with the miRNeasy Mini Kit (Qiagen), including DNA digestion and a single elution with RNase-free water. RNA quality was high, with an average RIN of 9.5, yielding approximately 3620 ng for alveolar macrophages (1E6 cells) and 6720 ng for monocyte-derived macrophages (2E6 cells). RNA sequencing for AM vs. MDM experiments were conducted using Clontech SMARTer stranded v2 Pico Mammalian kit. Sequencing was performed on an Illumina NovaSeq 6000 at Fred Hutchinson Cancer Research Center (Seattle, WA) and Vanderbilt University Medical Center (VANTAGE). Quality checks were performed using FastQC, with adapter removal by AdapterRemoval, alignment to the human genome (GRCh38) via STAR, and alignment filtering using Picard and samtools. Read counts were calculated with Rsubread, and all samples met quality standards.

## Acknowledgments

We thank the study participants whose invaluable contribution made this project possible. We appreciate the assistance of the bronchoscopy team at Harborview Medical Center in Seattle, WA. We also acknowledge the University of Washington sequencing core for expertise. We thank members of the Rothchild, Campo, and Hawn labs for helpful discussions.

## Data and materials availability

Raw and processed RNA-sequencing data can be accessed from the National Center for Biotechnology Information (NCBI) Gene Expression Omnibus (GEO) database under accession numbers GSE212203 (murine AM), GSE162620 (murine BMDM), and GSE236156 (human AM and MDM). All code is available on GitHub at https://github.com/hawn-lab/AM-MDM-cross-species

## FIGURE LEGENDS

**Figure S1.**
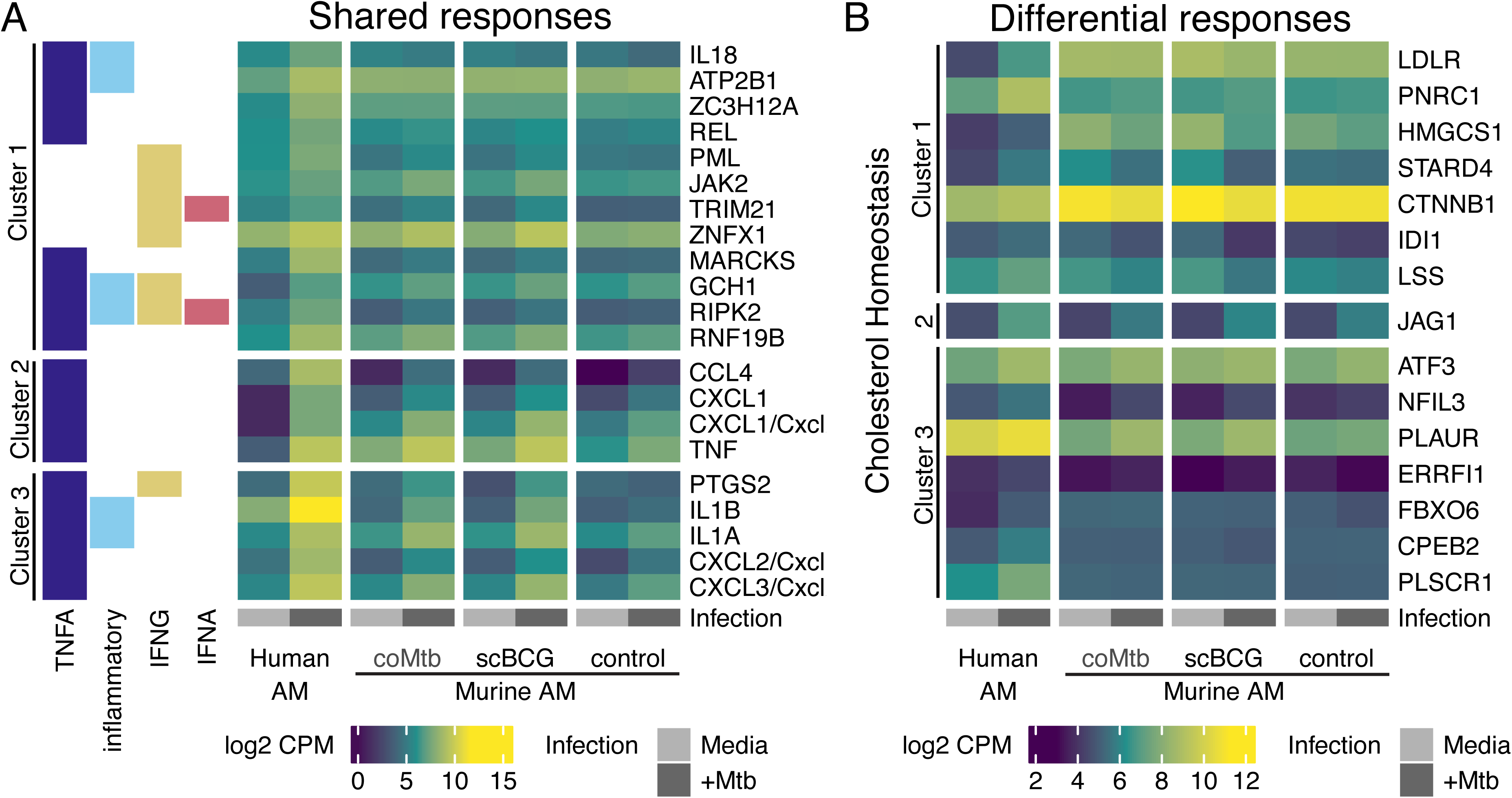
Gene expression in shared and unique human and murine AM responses to Mtb infection. Gene set enrichment analysis (GSEA) of Mtb vs media gene expression in human and murine AMs. (A) Shared responses were defined as gene sets significantly enriched in the same direction in human and murine AMs (FDR < 0.01). Genes were filtered to those expressed in both human and murine AMs, GSEA leading-edge in at least one selected pathway, and among the top 20 differentially expressed genes (DEGs) in any one AM (FDR < 0.1). Heatmap depicts mean media and +Mtb normalized log_2_ counts per million (CPM) expression in human and murine AMs. The same row ordering is applied from Figure 2B, and human HGNC symbols are used when human and murine gene names are identical. Left color annotation indicates the presence/absence of genes in each selected pathway including TNFA signaling via NK-kB (TNFA), inflammatory response (inflammatory), IFNG response (IFNG), and IFNA response (IFNA). (B) Unique responses were defined as gene sets significantly enriched in opposite directions in human and/or murine AM (FDR < 0.1). Genes were filtered and heatmaps were clustered similar to (A) but for the cholesterol homeostasis pathway. The same row ordering is applied from Figure 2D.

**Figure S2.**
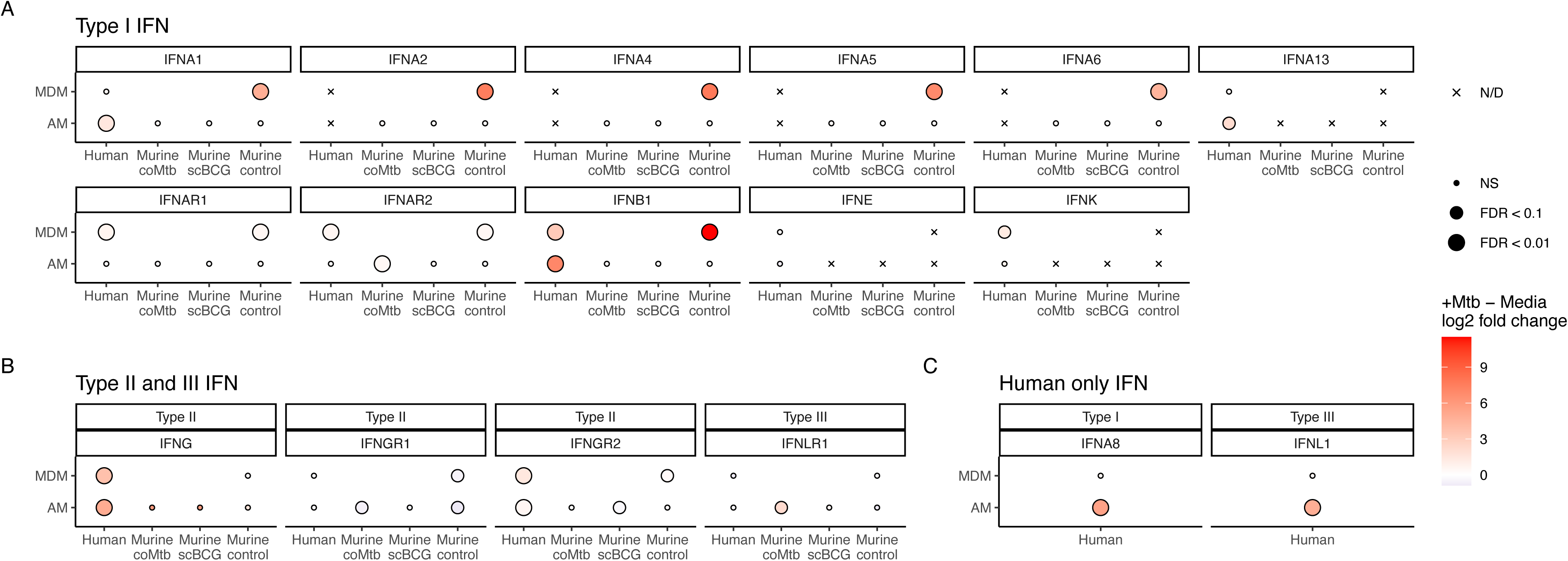
IFN expression in human and murine AM and MDM. Mtb-infected vs media fold change of normalized log_2_ counts per million (CPM) IFN gene expression. (A) Type I, (B) Type II, and Type III IFNs present in both human and murine genomes. (C) IFNs present in only the human transcriptome. No IFN were present in only the murine transcriptome. Color indicates log_2_ fold change (red increased expression with +Mtb) and size indicates FDR significance. Genes with expression below detection (N/D) are indicated by X. Expression was not measured in MDMs for the murine scBCG or coMtb models (blanks).

**Figure S3.**
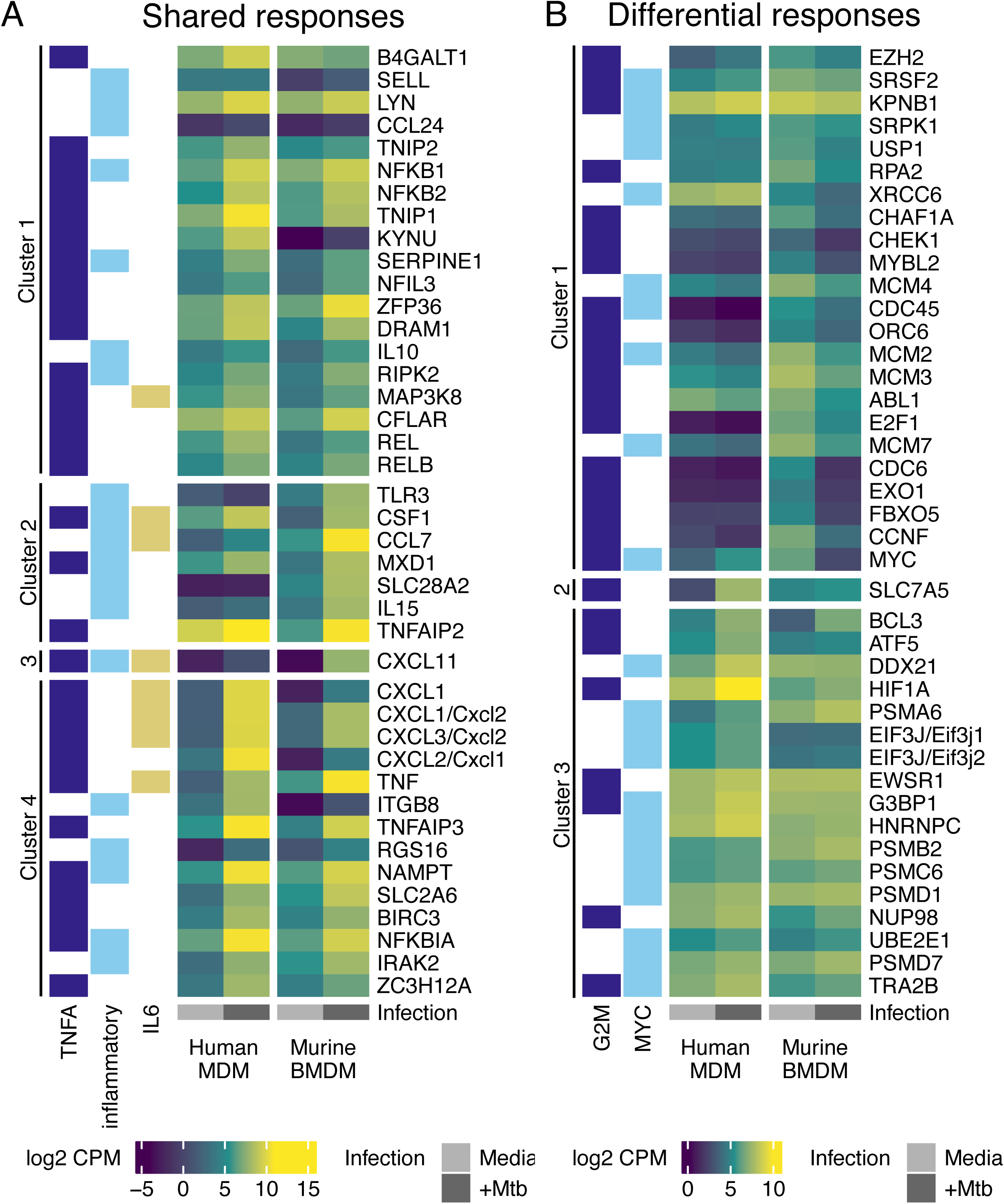
Gene expression in shared and unique human and murine MDM responses to Mtb infection. Gene set enrichment analysis (GSEA) of Mtb vs media gene expression in human and murine MDMs. (A) Shared responses were defined as gene sets significantly enriched in the same direction in human and murine MDMs (FDR < 0.01). Genes were filtered to those expressed in both human and murine MDMs, GSEA leading-edge in at least one selected pathway, and among the top 20 differentially expressed genes (DEGs) in any one MDM (FDR < 0.1). Heatmap depicts mean media and +Mtb normalized log_2_ counts per million (CPM) expression in human and murine MDMs. The same row ordering is applied from Figure 3B, and human HGNC symbols are used when human and murine gene names are identical. Left color annotation indicates the presence/absence of genes in each selected pathway including TNFA signaling via TNFA signaling via NK-kB (TNFA), inflammatory response (inflammatory), and IL6 JAK STAT3 signaling (IL6). (B) Unique responses were defined as gene sets significantly enriched in opposite directions in human and/or murine MDMs (FDR < 0.1). Genes were filtered and heatmaps were clustered similar to (A) but for G2M checkpoint (G2M) and MYC targets v1 (MYC) pathways. The same row ordering is applied from Figure 3D.

**Figure S4.**
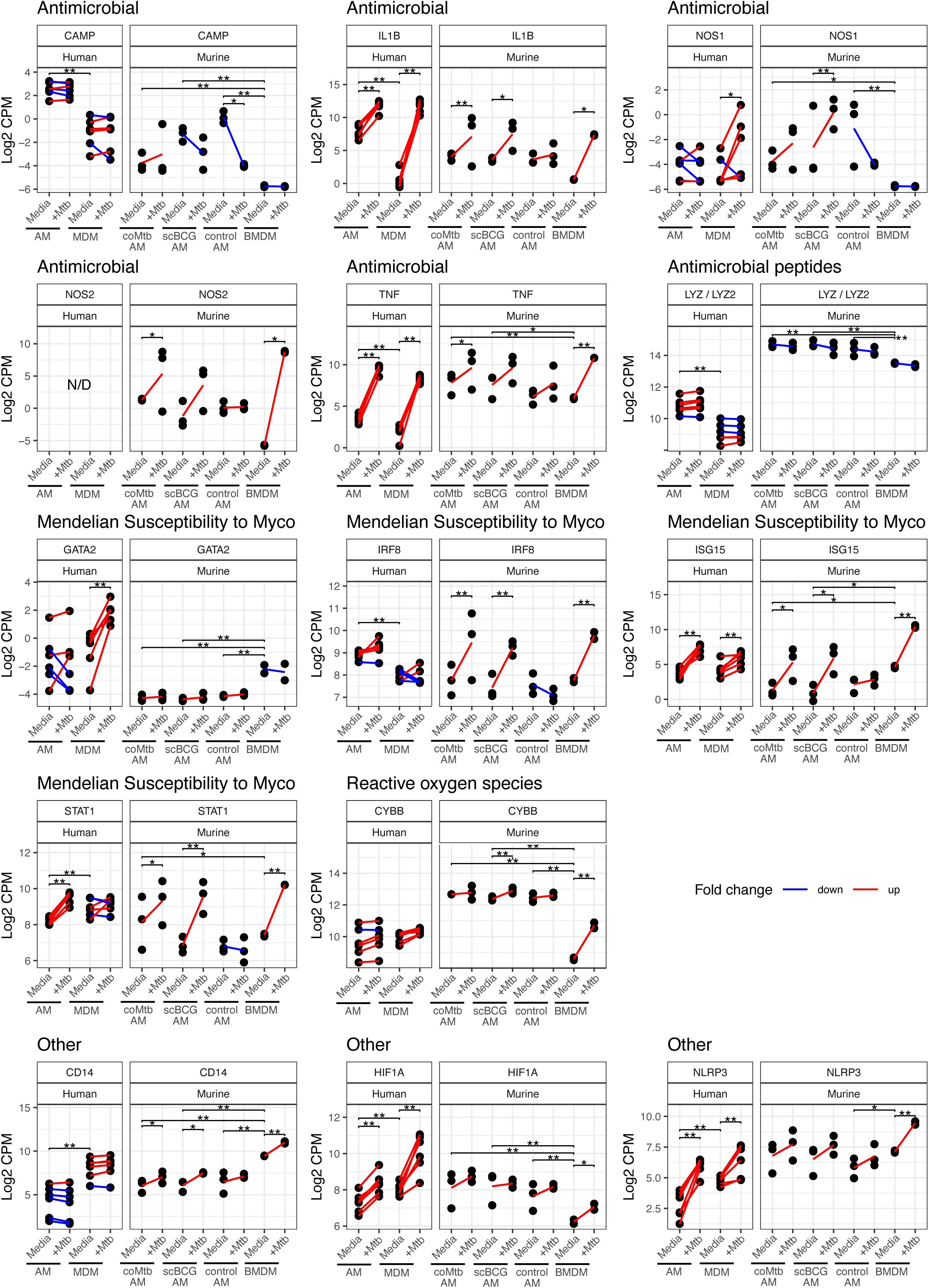
TB associated genes in human and murine AMs and MDMs. Log_2_ normalized counts per million (CPM) expression of select (A) antimicrobial, (B) antimicrobial peptides, (C) reactive oxygen species, (D) reactive nitrogen intermediates, (E) autophagy, (F) Mendelian susceptibility, and (G) other TB disease associated gene expression in human and murine AM and B/MDM. Paired media and +Mtb samples are connected by a line for human samples while a single line indicates the difference in mean media and +Mtb expression for unpaired murine samples. These lines are colored by positive (red) vs negative (blue) fold change. Significance is indicated for Mtb infection within each cell type and between cell types in the media condition. ** FDR < 0.01; * FDR < 0.1 Myco: Mycobacterium, N/D: not detected.

**Table S1. Gene-level linear model.** Human AM and MDMs were modeled for Mtb infection with a random effect for donor, ∼cell*Mtb + 1|donor. Murine AM and BMDMs were modeled for Mtb infection vaccination (control, coMtb, scBCG), ∼cell*Mtb. Each tab includes results for one cell type and contrast for Mtb vs media or cell types within media. estimate: log_2_ fold change estimate, pval: p-value, FDR: Benjamini-Hochberg false discovery rate corrected p-value

**Table S2. Gene set enrichment analysis (GSEA).** Mtb-infected minus media linear model estimates were used in GSEA of the MSigDB Hallmark gene set database. Each tab includes results for one cell type. NES: normalized enrichment score, pval: p-value, FDR: Benjamini-Hochberg false discovery rate corrected p-value.

